# MRP1 inhibition by lipid-derived electrophiles during ferroptosis illustrates a role for protein alkylation in ferroptotic cell death

**DOI:** 10.1101/2023.10.09.559028

**Authors:** Antonius T. M. Van Kessel, Gonzalo Cosa

## Abstract

Ferroptosis is a regulated form of cell death characterized by lipid peroxidation and lipid hydroperoxide (LOOH) generation that offers new therapeutic opportunities. However, the molecular mechanism through which LOOH accumulation leads to cell death remains poorly understood. Importantly, LOOH breakdown forms truncated phospholipids (PLs) and highly reactive lipid-derived electrophiles (LDEs) capable of altering protein function through cysteine alkylation. While truncated PLs have been shown to mediate ferroptotic membrane permeabilization, a functional role for LDEs in the ferroptotic cell death mechanism has not been established. Here, using multidrug resistance protein 1 (MRP1) activity as an example, we demonstrate that LDEs mediate altered protein function *during ferroptosis*. Applying live cell fluorescence imaging, we first identified that inhibition of MRP1-mediated LDE detoxification occurs across a panel of ferroptosis inducers (FINs) with differing mechanisms of ferroptosis induction (Types I-IV FINs erastin, RSL3, FIN56 and FINO_2_). This MRP1 inhibition was recreated by both initiation of lipid peroxidation and treatment with the LDE 4-hydroxy-2-nonenal (4-HNE). Importantly, treatment with radical-trapping antioxidants prevented impaired MRP1 activity when working with both FINs and lipid peroxidation initiators but not 4-HNE, pinpointing LDEs as the cause of inhibited MRP1 activity during ferroptosis. Our findings, when combined with reports of widespread LDE-alkylation of key proteins during ferroptosis, sets a precedent for LDEs as critical mediators of ferroptotic cell death. LOOH breakdown to truncated phospholipids and LDEs may fully explain membrane permeabilization and modified protein function during late stage ferroptosis, offering a unified explanation of the molecular ferroptotic cell death mechanism.

## Introduction

Ferroptosis is a regulated cell death mechanism defined by elevated levels of iron-dependent lipid peroxidation (1–4). The potential of ferroptosis induction as a cancer therapy (5–7) and the association of ferroptosis with numerous diseased states, including ischemia reperfusion injury (8, 9) and neurodegenerative disease (10, 11), has firmly established the importance of this mode of cell death (12, 13). The central mechanism of ferroptosis involves radical chain-mediated autoxidation (lipid peroxidation) of polyunsaturated fatty acid containing phospholipids (PUFA-PLs) to form high levels of cellular lipid hydroperoxides (PUFA-OOH, **Figure 1A**) (3, 14). A complex network of cellular antioxidant, iron and lipid metabolism mechanisms are involved in regulating cellular susceptibility to lipid peroxidation, and by extension ferroptosis (4, 12, 15). This current understanding has been shaped by the identification of small molecule ferroptosis inducers (FINs, see **Figure 1A** for a subset of these molecules and how they impinge on lipid peroxidation) and inhibitors of ferroptosis (16–19).

**Figure 1.**
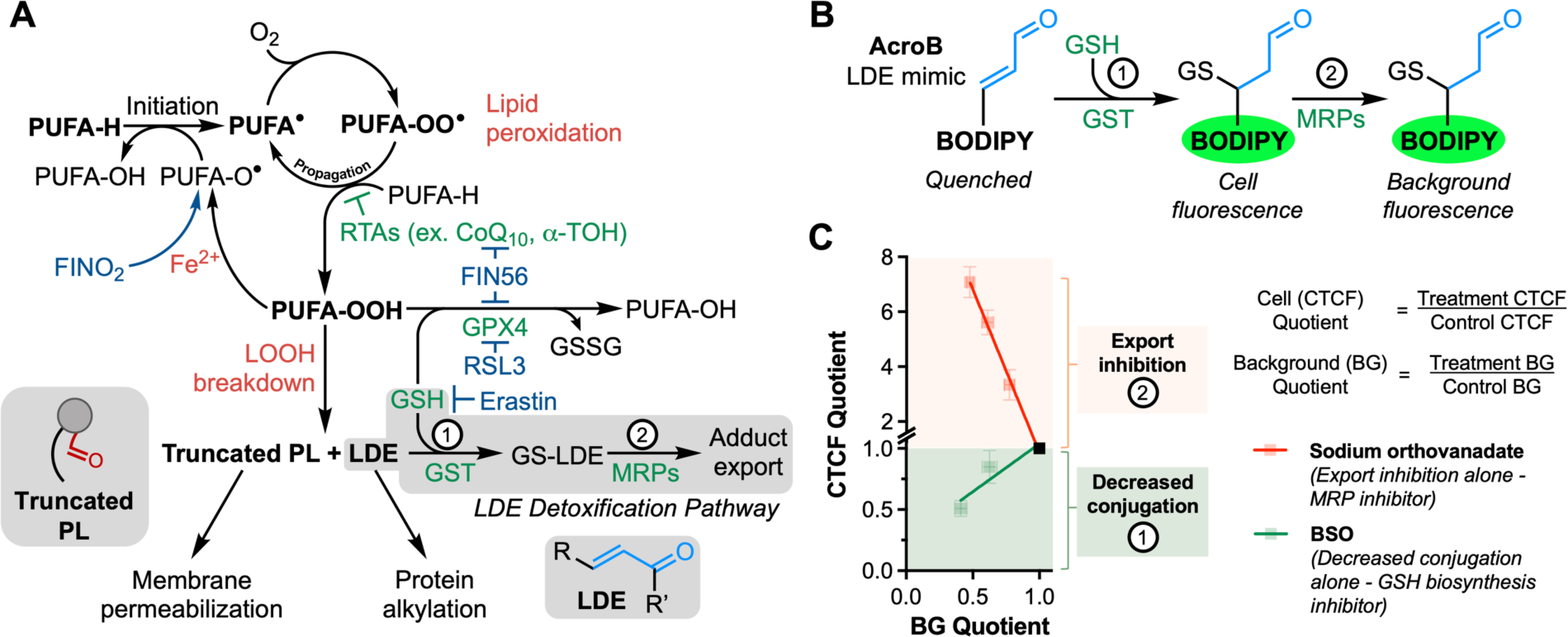
Evaluation of lipid-derived electrophile (LDE) detoxification during ferroptosis using ElectrophileQ. **A**, Scheme of the integrated nature of ferroptosis and LDE metabolism. Induction of ferroptosis can be achieved with a variety of small molecules that, through different mechanisms, lead to increased levels of lipid peroxidation. In the presence of iron, FINO_2_ increases initiation rates of lipid peroxidation. RSL3 inhibits GPX4, the enzyme responsible for reduction of lipid hydroperoxides to lipid-derived alcohols. FIN56 treatment leads to depletion of CoQ_10_ and GPX4 degradation. Erastin prevents GSH biosynthesis through inhibition of cystine uptake. LOOHs break down to form truncated PL and LDEs. The two-step LDE detoxification pathway involves i) LDE conjugation with GSH catalyzed by GST and ii) export of the formed GS-LDE adduct through MRP channels. **B**, The fluorogenic (turn-on) LDE-mimic AcroB enables study of LDE metabolism as levels of cell fluorescence and background fluorescence depend on both the conjugation and export steps of the LDE detoxification pathway. **C**, The ElectrophileQ assay in HT1080 cells. Comparing the level of corrected total cell fluorescence (CTCF) and background (BG) fluorescence in test conditions to control values reports on the level of AcroB conjugation and export. Plotting the CTCF quotient vs. the BG quotient for the glutathione inhibitor BSO establishes a standard curve for decreased conjugation alone (lower cellular GSH decreases AcroB conjugation, decreasing CTCF and BG relative to control). The ATPase inhibitor sodium orthovanadate establishes a standard curve for export inhibition alone (prevention of adduct export increases CTCF and decreases BG relative to control). Standard curve data reproduced from reference 36. The control point is presented as a visual aid at 1.0-1.0. Any points between the standard curves signal a combination of both modes of LDE detoxification inhibition, with increasing contribution of decreased conjugation as CTCF quotient decreases.

Mapping the steps of ferroptosis between lipid hydroperoxide accumulation and eventual membrane permeabilization and cell lysis is an active area of research (20, 21). Following lipid peroxidation, breakdown of the inherently unstable PUFA-OOHs has been shown to generate truncated phospholipids (PLs) and diffusible lipid-derived electrophiles (LDEs, **Figure 1A**) such as 4-hydroxy-2-nonenal (4-HNE) and malondialdehyde (MDA) (22). The formed truncated PLs, which are known to permeabilize lipid membranes (23–26), are promising candidates to cause loss of membrane integrity and mediate ferroptotic membrane permeabilization (27). Consistent with this model, during late-stage ferroptosis, it has been reported that plasma membrane pores form leading to cell swelling, calcium influx and recruitment of membrane repair machinery (28–31). Recently, activation of mechanosensitive Piezo1 and TRP channels and inactivation of the sodium-potassium (Na^+^/K^+^)-ATPase were identified as critical steps contributing to loss of the plasma membrane cation gradient (32).

Although elevated LDE levels are well accepted as a ferroptosis marker (33–35), and their impaired detoxification constitutes a hallmark of ferroptosis (36), a functional role for these highly reactive species has not been well evaluated in the ferroptosis context. LDEs are α,β-unsaturated aldehydes or ketones that react primarily with cysteine nucleophiles in the cell (**Figure 1A**) (37). LDE alkylation of protein thiols can modify enzyme catalytic activity, protein structure, and membrane channel properties, where widespread LDE alkylation can significantly impair cell function (38). The critical role of cysteine (or selenocysteine) residues in proteins involved in redox metabolism/ferroptosis regulation (ex. FSP1, iPLA2β, GPX4) positions LDEs as critical mediators of cellular damage downstream of PUFA-OOH during ferroptosis. In fact, a screen of LDE alkylation during ferroptosis identified over 1000 modified proteins (39). While LDEs hold great promise as critical mediators of cell damage, a demonstration of LDE-mediated altered protein function *during ferroptosis* is currently missing.

Here, we establish a connection between LDE-alkylation and modified protein function during ferroptosis, illustrating the ramifications of PUFA-OOH generation and subsequent breakdown through providing a putative molecular mechanism of ferroptotic cell death. We chose multidrug resistance protein 1 (MRP1) activity as a test case. MRPs are responsible for the export of glutathione (GSH)-LDE adducts in the LDE detoxification pathway (**Figure 1A**) (40, 41). MRP1 has independently been shown to be alkylated during ferroptosis (39) and inhibited by LDEs (42–44). Applying a fluorescence-based LDE detoxification assay we developed, ElectrophileQ (36), here we perform a comprehensive evaluation of LDE detoxification impairment during ferroptosis. Through testing a series of canonical ferroptosis inducers, we show that inhibition of MRP1-mediated LDE-adduct export occurs across all tested modes of ferroptosis induction regardless of the mechanism (i.e., Type I-IV FINs). Next, we demonstrate that this LDE-adduct export inhibition is recapitulated by the addition of lipid peroxidation initiators and exogenous LDEs. However, while radical trapping antioxidant (RTA) treatment during FIN or lipid peroxidation exposure prevents inhibition of MRP1 function, RTA treatment during exposure to exogenous LDEs is insufficient to prevent LDE-adduct export inhibition. Together, our results indicate that LDEs generated during ferroptosis inhibit MRP1 channels. Given the widespread LDE-mediated protein alkylation mapped in ferroptotic cells, this link between LDEs and modified MRP1 function establishes a precedent for LDEs as mediators of altered cell metabolism and advances our understanding of ferroptotic cell death. Our results position LDEs alongside truncated PLs as the critical mediators of cellular damage during ferroptosis.

## Results

To evaluate LDE detoxification in live cells we applied the ElectrophileQ assay (36) which employs a fluorogenic (turn-on) LDE mimic, AcroB, as a reporter (**Figure 1B**) (45). LDE detoxification in healthy cells has two steps: i) LDE conjugation with glutathione (GSH) catalyzed by the glutathione *S*-transferase (GST) class of enzymes and ii) active export of the formed GS-LDE adducts through MRP channels (**Figure 1A**) (40, 41, 46). AcroB facilitates visualization of the LDE detoxification pathway in real time as first the probe becomes fluorescent following reaction with GSH and next the resulting fluorescent adducts are exported from the cell through MRP channels leading to increased background fluorescence (**Figure 1B**). AcroB can thus readily report on MRP channel activity during ferroptosis.

Using widefield fluorescence microscopy, comparison of AcroB cellular and background fluorescence to control values generated the ElectrophileQ plot, providing a functional readout of cellular ability to execute both steps of LDE detoxification (**Figure 1C**). To interrogate the mechanism of LDE detoxification impairment in test conditions, (i) sublethal L-buthionine sulfoximine (BSO) and (ii) sodium orthovanadate are used to generate standard curves for inhibition of only the first or second steps of the LDE detoxification pathway, respectively (**Figure 1C**). As a GSH biosynthesis inhibitor, the BSO line on the ElectrophileQ plot represents the relationship between the change in AcroB cell fluorescence (CTCF – corrected total cell fluorescence) and background fluorescence (BG) relative to control conditions when only the first step of LDE detoxification is impaired. Likewise, the line generated by the ATPase inhibitor sodium orthovanadate represents the relationship between CTCF and BG quotients relative to control conditions when only MRP-mediated LDE-adduct export is impaired (**Figure 1C**) (36). Since sodium orthovanadate is a promiscuous inhibitor of membrane ATPase channels, we tested two additional, more selective, MRP1 inhibitors (Reversan and MK571, **Figure S1**) (47, 48). The ElectrophileQ response of both Reversan and MK571 closely matched the sodium orthovanadate line, confirming this as a suitable standard for MRP1 inhibition on the ElectrophileQ plot (**Figure S1**).

### RSL3, FINO_2_ and FIN56 induce LDE-adduct export impairment

To evaluate if impairment of LDE-adduct export occurs during all forms of ferroptosis induction, we initially applied the ElectrophileQ assay to screen LDE detoxification ability across a series of ferroptosis inducers (FINs). Small molecule inducers of ferroptosis increase lipid peroxidation through multiple mechanisms and FINs are often divided into four classes (**Figure 1A**). Type I FINs, such as erastin, lead to glutathione (GSH) depletion and indirect inhibition of GPX4, the GSH-dependent enzyme responsible for the reduction of PUFA-OOH to less reactive PUFA-OH (**Figure 1A**). Type II FINs include RSL3 and directly inhibit GPX4 activity (16). Type III FINs lead to degradation of GPX4 and depletion of cellular radical-trapping antioxidants (RTAs) with the prototypical example being FIN56, which depletes cellular coenzyme Q_10_ (CoQ_10_) (18). In the presence of iron, the Type IV FIN FINO_2_ increases the rate of lipid peroxidation initiation (17).

We initially tested Types II-IV FINs, specifically FINO_2_, FIN56 and a relatively low [RSL3] to match the timescale of ferroptosis induction by the other two compounds (**Figure 2**). Visually, treatment of HT1080 cells with all three FINs led to fluorescent GS-AcroB adduct retention in the cytoplasm compared to control conditions (**Figure 2A**, additional images in **Figure S2A-C**). Fluorescence quantification of these images showed an incubation-time dependent increase in CTCF and decrease in BG fluorescence across all 3 FINs compared to control conditions (**Figure 2B-D**, middle 2 time trajectory panels). Evaluating these changes using ElectrophileQ analysis mapped LDE detoxification impairment in close agreement with the sodium orthovanadate line, demonstrating that RSL3, FINO_2_ and FIN56 all induce LDE-adduct export impairment (**Figure 2B-D**, right panel). Slight deviations from the sodium orthovanadate line for all three FINs, specifically at the longest incubation times presented, suggests that cells in later stages of ferroptosis induced by these compounds additionally experience impairment of LDE conjugation ability, potentially through decreased GST activity. The LDE detoxification impairment induced by FINO_2_ and FIN56 was completely prevented by pre-treatment with the radical-trapping antioxidant (RTA) phenoxazine (PHOXN, **Figure S2D**), similar to RSL3 as we previously demonstrated (36).

**Figure 2.**
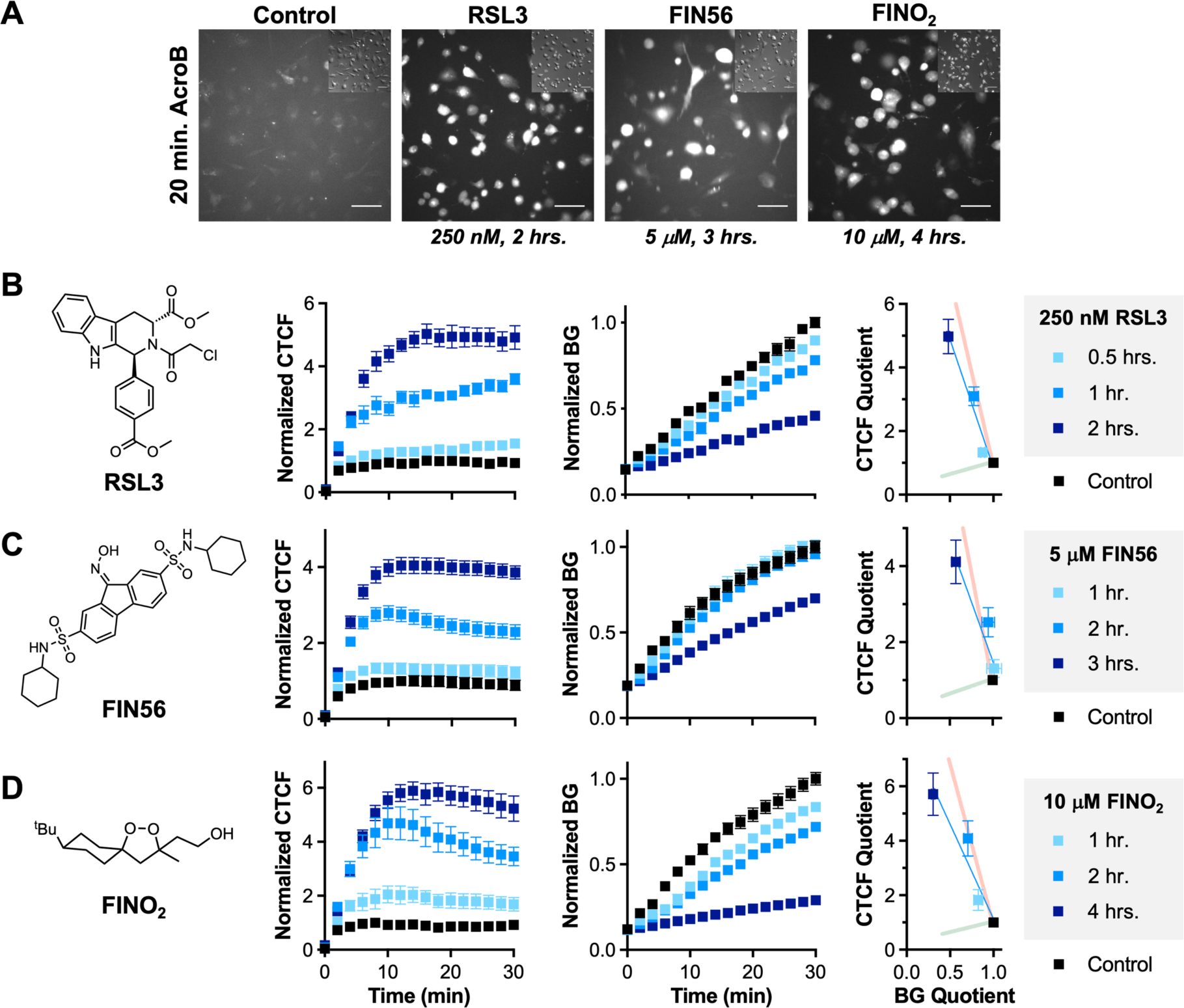
Evaluation of LDE detoxification following treatments with RSL3, FIN56 and FINO_2_ reveals consistent LDE adduct export impairment. **A**, Representative 20*×* AcroB widefield fluorescence and DIC (inset) images of HT1080 cells following treatment with RSL3, FIN56 or FINO_2_ show consistent increase in cell fluorescence and decrease in background fluorescence compared to control. Images were taken every 2 minutes for 30 minutes following application with 100 nM AcroB (image 20 minutes following AcroB application is shown). Image is presented for the longest incubation performed with each FIN (incubation with FINs performed before AcroB application) – for images of additional treatments see **Figure S2**. Scale bar is 64 μm. α_ex_ = 488 nm (0.1 mW). **B-D**, Chemical structures of FINs utilized, normalized average CTCF, normalized average BG, and ElectrophileQ plots following indicated treatment times and concentrations for (**B**) RSL3, (**C**) FIN56 and (**D**) FINO_2_ obtained from 20*×* images. AcroB was added following the indicated incubation time with each FIN. Each data set was normalized to maximum CTCF and BG values obtained using controls performed in parallel. The ElectrophileQ plot analyzes the CTCF and BG values calculated for the images obtained 20 minutes following AcroB application. Sodium orthovanadate and BSO standard curves presented as visual aids. All 3 FINs produce relationships on the ElectrophileQ plot that reflect LDE detoxification impairment dominated by LDE-adduct export impairment. Presented values are average of minimum n = 4 field of view (FOV) for all conditions ± SEM.

Despite differing mechanisms of ferroptosis induction, the Types II-IV FINs RSL3, FIN56 and FINO_2_ all lead to LDE detoxification impairment through a consistent mechanism characterized by inhibition of MRP1-mediated LDE-adduct export.

### Erastin induces LDE conjugation and adduct export impairment

We next tested the Type I FIN erastin, which inhibits system x_c_^-^ preventing the import of the GSH precursor cystine (19). Erastin is also known to bind to VDAC channels and increase mitochondrial reactive oxygen species (ROS) generation (49–51). Here, we applied the ElectrophileQ method to evaluate LDE detoxification during erastin-induced ferroptosis (**Figure 3**). Given GSH depletion will directly impact the conjugation step of LDE detoxification, we anticipated erastin would correlate well with BSO on the ElectrophileQ plot (i.e., decrease in both CTCF and BG). However, while overall AcroB fluorescence did visually decrease following erastin treatment for 10-11 hrs. (**Figure 3A**), retention of AcroB adducts were observed and quantification demonstrated an increase in CTCF and decrease in BG (**Figure 3B**, middle 2 panels), placing this treatment between the sodium orthovanadate and BSO lines (**Figure 3B**, right panel). This pattern signals inhibition of GSH conjugation but, importantly, also indicates that LDE-adduct export impairment does occur during treatment with the Type I FIN erastin.

**Figure 3.**
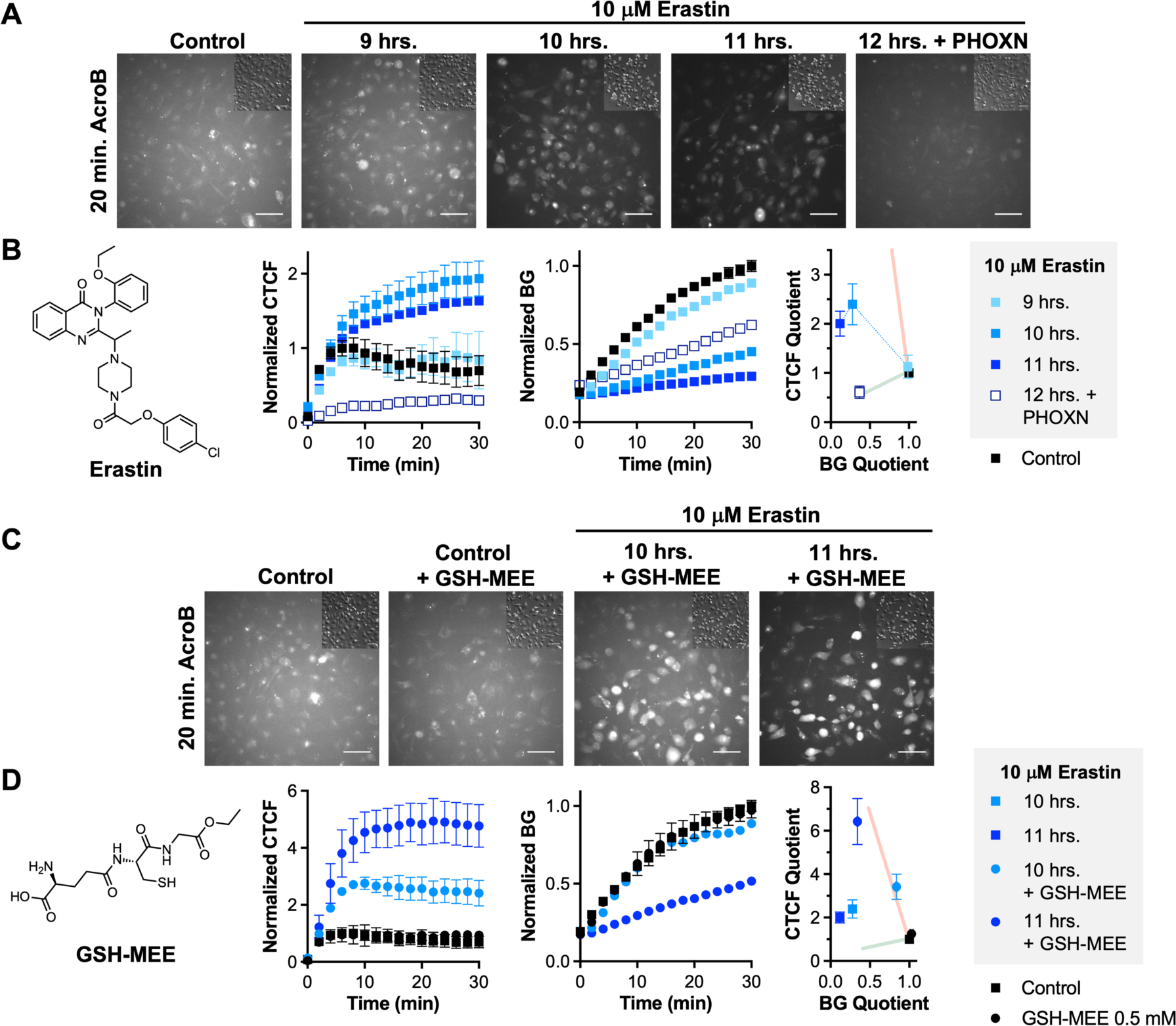
Evaluation of LDE detoxification following erastin treatment reveals mixed GSH conjugation impairment and LDE-adduct export impairment. **A**, Representative 20× AcroB widefield fluorescence and DIC (inset) images of HT1080 cells following treatment with 10 μM erastin for 9-11 hrs. and following 1 μM PHOXN and 12 hr. 10 μM erastin treatment compared to control conditions. **B**, Chemical structure of erastin, normalized average CTCF, normalized average BG, and ElectrophileQ plots for conditions presented in (**A**). 9-11 hrs. erastin incubation produced LDE detoxification impairment consistent with a mix of GSH conjugation impairment and LDE-adduct export impairment. Line connecting 9, 10 and 11 hr. erastin treatment points is a visual aid. PHOXN + 12 hrs. erastin incubation produced LDE detoxification impairment consistent with GSH conjugation impairment alone. **C**, Representative 20× AcroB widefield and DIC (inset) images of HT1080 cells following no treatment (Control) or treatment with 10 μM erastin for 10-11 hrs. followed by 30-minute 0.5 mM GSH-MEE treatment prior to AcroB application. **D**, Chemical structure of GSH-MEE, normalized average CTCF and normalized average BG plots for conditions presented in (**C**) and ElectrophileQ plots for selected conditions presented in (**A**) and (**C**). Re-introduction of GSH through GSH-MEE treatment improves visualization of the LDE-adduct export impairment caused by erastin treatment. Images were taken every 2 minutes for 30 minutes following application with 100 nM AcroB (image 20 minutes following AcroB application shown). α_ex_ = 488 nm (0.1 mW). Scale bar is 64 μm. AcroB added following the indicated incubation time with erastin and/or GSH-MEE. Each data set was normalized to maximum CTCF and BG values obtained using controls performed in parallel. The ElectrophileQ plot analyzes the CTCF and BG values calculated for the images obtained 20 minutes following AcroB application. Sodium orthovanadate and BSO standard curves presented as visual aids. Presented values are average of minimum n = 4 field of view (FOV) for all conditions ± SEM.

To test if erastin-induced impairment of MRP1-mediated LDE-adduct export is dependent on lipid peroxidation, we treated HT1080 cells with PHOXN prior to erastin exposure. While 12 hr. erastin incubation led to cell death, pre-incubation with PHOXN restored cell viability (**Figure 3A**, far right panel). ElectrophileQ analysis of the PHOXN + 12 hr. erastin condition showed that the measured CTCF and BG quotients matched well with the BSO line, consistent with inhibition of GSH conjugation alone and no LDE adduct export impairment (**Figure 3B**), demonstrating that the LDE-adduct export inhibition caused by erastin is dependent on lipid peroxidation.

The mixed inhibition of LDE detoxification following erastin treatment makes it difficult to compare the level of LDE-adduct export impairment during erastin-induced ferroptosis to that caused by RSL3, FINO_2_ and FIN56 treatment. To better resolve the inhibition of LDE-adduct export following erastin treatment, we re-introduced GSH to HT1080 cells following 10-11 hrs. of erastin exposure using treatment with 0.5 mM GSH-monoethyl ester (GSH-MEE, **Figure 3C-D**). While this addition of GSH-MEE had no effect on AcroB signal in the absence of erastin, GSH-MEE application following erastin treatment produced high levels of AcroB cellular fluorescence reminiscent of that observed with RSL3, FINO_2_ and FIN56 (**Figure 3C**). Quantification of CTCF and BG along with ElectrophileQ analysis (**Figure 3D**) showed good agreement between erastin + GSH-MEE and the sodium orthovanadate line, demonstrating that the LDE-adduct export impairment caused by erastin is consistent with inhibition of MRP activity and marked by a similar level of LDE-adduct export impairment as observed with RSL3, FINO_2_ and FIN56-induced ferroptosis. All tested forms of ferroptosis induction led to inhibition of MRP1-mediated LDE-adduct export.

The characterization of LDE detoxification during erastin-induced ferroptosis suggests that erastin activity aside from just GSH depletion contributes to cell death. This is supported by the fact that erastin causes a significantly different response on the ElectrophileQ plot than BSO (**Figure 3**) (36). These data support that alternative mechanisms of action of erastin, including the opening of VDAC channels and subsequent increased mitochondrial ROS generation following mitochondrial hyperpolarization, are important during erastin-induced ferroptosis (49, 50, 52).

### Induction of lipid peroxidation is sufficient to inhibit LDE-adduct export

Following the identification of LDE-adduct export impairment across all tested FINs (Type I-IV), we aimed to evaluate the minimum requirements for induction of this cellular state and determine the mechanism of MRP1 inhibition during ferroptosis.

In order to examine if simply the induction of a cell death pathway impairs cellular ability to detoxify LDEs, we induced apoptosis in HeLa cells with etoposide (**Figure S3A-C**). Since etoposide inhibits cell division, the required prolonged treatment led to cell growth and increase in CTCF (**Figure S3B**). However, LDE detoxification impairment during apoptosis was ruled out through correction for this cell growth, as AcroB signal per cell pixel is unchanged compared to the control (**Figure S3C**). The induction of apoptosis does not cause LDE detoxification impairment.

To evaluate if the induction of general ROS-mediated oxidative stress induces LDE-detoxification impairment, we treated cells with hydrogen peroxide (H_2_O_2_). Treatment of HeLa cells with 2 mM H_2_O_2_ for 2 hrs. induced significant morphological changes (**Figure S3D**), cellular retention of GS-AcroB adducts (**Figure 4A**, quantification in **Figure S3E**) and a response on the ElectrophileQ plot matching the LDE-adduct export impairment observed during ferroptosis (**Figure 4B**). Notably, while treatment with the RTA PHOXN did not lead to recovery of control cell morphology (**Figure S3D**), PHOXN pre-treatment did prevent the H_2_O_2_-induced AcroB CTCF and BG response (**Figure S3E**) and promoted recovery of LDE detoxification ability as shown by the ElectrophileQ plot (**Figure 4B**). The lipophilic RTA PHOXN restoring LDE detoxification during H_2_O_2_ treatment further suggests that lipid peroxidation, and not simply cellular ROS generation, is required for the LDE-adduct export impairment during ferroptosis. The induction of ferroptosis-like LDE detoxification impairment by H_2_O_2_ is of relevance given the critical role enzymes that generate H_2_O_2_ (ex. NOX4) play in the initiation of lipid peroxidation by hydroxyl radicals during ferroptosis (53–55).

**Figure 4.**
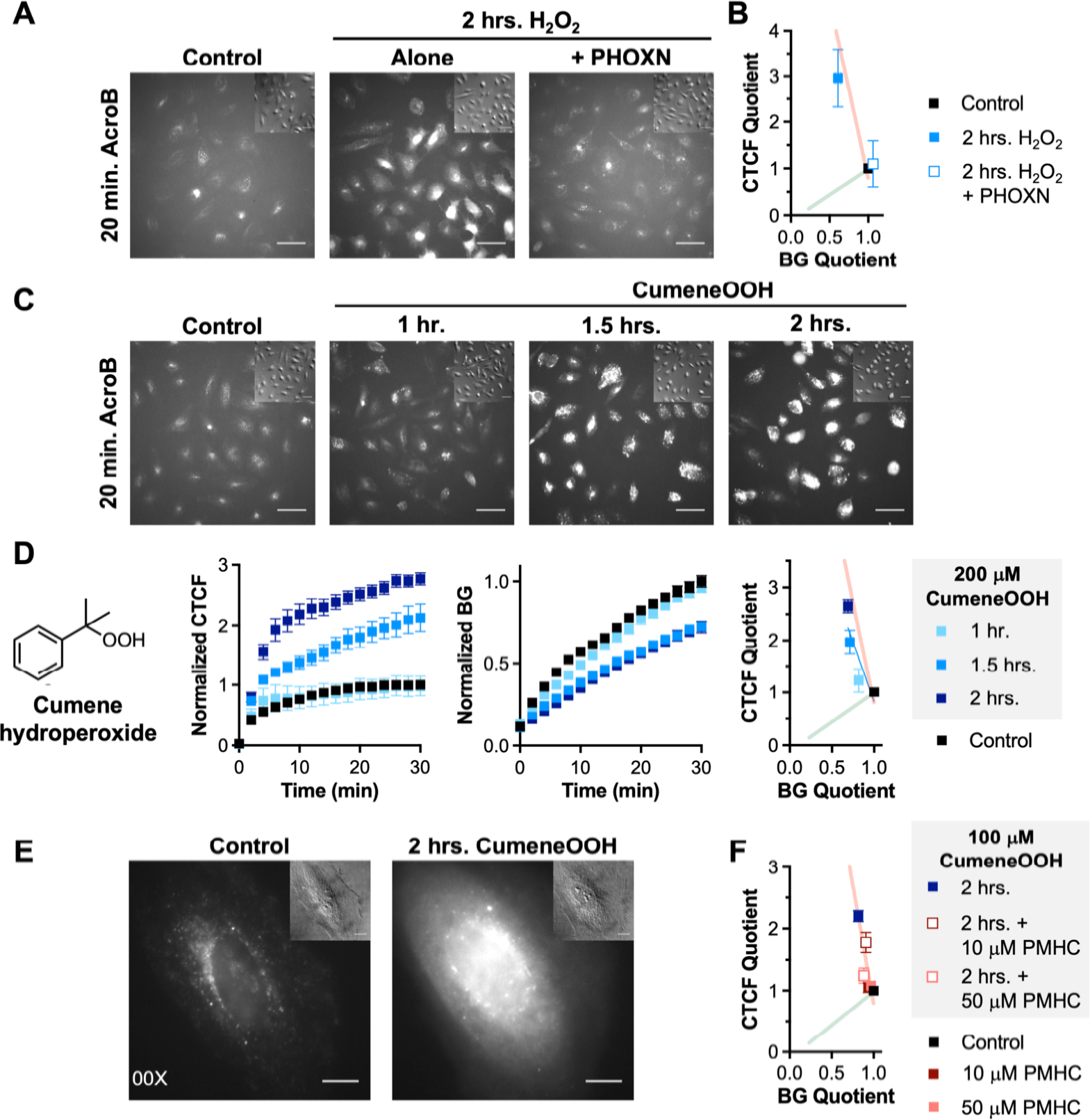
Induction of lipid peroxidation recapitulates the LDE detoxification impairment observed during ferroptosis. **A**, Representative 20*×* AcroB widefield fluorescence and DIC (inset) images of HeLa cells following 2 mM H_2_O_2_ treatment with/without 1 μM PHOXN pre-treatment compared to control conditions. **B**, ElectrophileQ plot for conditions presented in (**A**). Corresponding normalized average CTCF and BG plots and additional images presented in **Figure S3D-E**. H_2_O_2_ treatment produced an ElectrophileQ response consistent with LDE adduct export impairment that was recovered by PHOXN pre-treatment. **C**, Representative 20*×* AcroB widefield and DIC (inset) images of HeLa cells following 1-2 hr. treatment with 200 μM cumene hydroperoxide (cumeneOOH) and 20 μM copper (II) sulfate compared to control conditions. **D**, Structure of the lipid peroxidation inducer cumene hydroperoxide, normalized average CTCF, normalized average BG, and ElectrophileQ plots for conditions presented in (**C**). ElectrophileQ plot shows cumeneOOH produces LDE detoxification impairment consistent with that obtained in ferroptosis. **E**, 100*×* HeLa cell AcroB fluorescence and DIC (inset) images after no treatment or 2-hour treatment with 200 μM cumeneOOH and 20 μM copper (II) sulfate and 20-minute application of AcroB. Images show retention of cellular AcroB adducts following cumeneOOH treatment compared to control conditions. **F**, ElectrophileQ plot for conditions where cumeneOOH treatment was preceded by PMHC treatment presented in **Figure S3F-G**. Antioxidant treatment recovered control LDE detoxification response in a dose-dependent manner. Images taken every 2 min. for 30 min. (20*×*) or every 30 sec. for 30 min. (100*×*) – images taken 20 minutes after AcroB application shown. α_ex_ = 488 nm (0.1 mW for 20*×*, 0.05 mW for 100*×*). Scale bar is 64 μm for 20*×* and 12 μm for 100*×*. AcroB added following the indicated treatment(s). Each data set was normalized to maximum CTCF and BG values obtained using controls performed in parallel. The ElectrophileQ plot analyzes the CTCF and BG values calculated for the images obtained 20 minutes following AcroB application. Sodium orthovanadate and BSO standard curves presented as visual aids. Presented values are average of minimum n = 4 field of view (FOV) for all conditions ± SEM.

To confirm that the induction of lipid peroxidation is sufficient to recapitulate the LDE-adduct detoxification impairment that occurs during ferroptosis, we induced lipid peroxidation in HeLa cells with cumene hydroperoxide (cumeneOOH, **Figure 4D**) and copper (II) sulfate treatment (**Figure 4C-F**) (56). Induction of lipid peroxidation in HeLa cells produced an incubation-time dependent increase in cytoplasmic retention of AcroB adducts (**Figure 4C-E**). The relationship between the increase in CTCF and decrease in BG as shown by the ElectrophileQ plot mimics that obtained during ferroptosis induction (**Figure 4D**, right panel). Treatment with the RTA 2,2,5,7,8-pentamethyl-6-chromanol (PMHC) recovered the ElectrophileQ signal towards the control response (**Figure 4F**, images and quantification in **Figure S3F-G**).

Together, these results support that increased levels of lipid peroxidation during ferroptosis are responsible for producing species that inhibit MRP-mediated LDE-adduct export.

### Electrophiles induce inhibition of MRP1-mediated LDE-adduct export

Although LDEs have been shown to alkylate many proteins during ferroptosis (39), a characterization of the functional implications of this high level of protein alkylation is lacking. Given that MRP1 (ABCC1) was identified in the list of proteins alkylated by LDEs during ferroptosis (39), and knowing that LDE alkylation can inhibit MRP1 function (42–44), we next explored whether alkylation of MRP1 by electrophiles accounts for the ferroptotic LDE-adduct export inhibition we have characterized.

To test if LDEs mediate inhibition of LDE-adduct export, we assessed LDE detoxification ability following treatment with a series of α,β-unsaturated carbonyl electrophiles (**Figure 5A**). The initial compound tested was N-ethyl maleimide (NEM, **Figure 5A**), a reagent widely used to deplete GSH and as a cysteine-specific alkylation reagent (57). Consistent with NEM-mediated GSH depletion, treatment of HeLa cells with 100 μM NEM prior to AcroB exposure greatly reduced CTCF and BG (**Figure S4A-B**) and produced an ElectrophileQ response that agrees well with the BSO line (**Figure 5C**). To test if GSH depletion may be masking alkylation-based inhibition of MRP channels by NEM, we supplemented NEM-treated cells with GSH-MEE. Cells treated with NEM followed by GSH-MEE showed the retention of AcroB adduct fluorescence characteristic of impaired LDE-adduct export (**Figure 5B**), a much greater increase in CTCF than BG relative to NEM alone (**Figure S4A-B**) and a mixed inhibition based on the ElectrophileQ response (**Figure 5C**). Therefore, treatment with the α,β-unsaturated carbonyl NEM leads to functional inhibition of MRP1-mediated LDE-adduct export.

**Figure 5.**
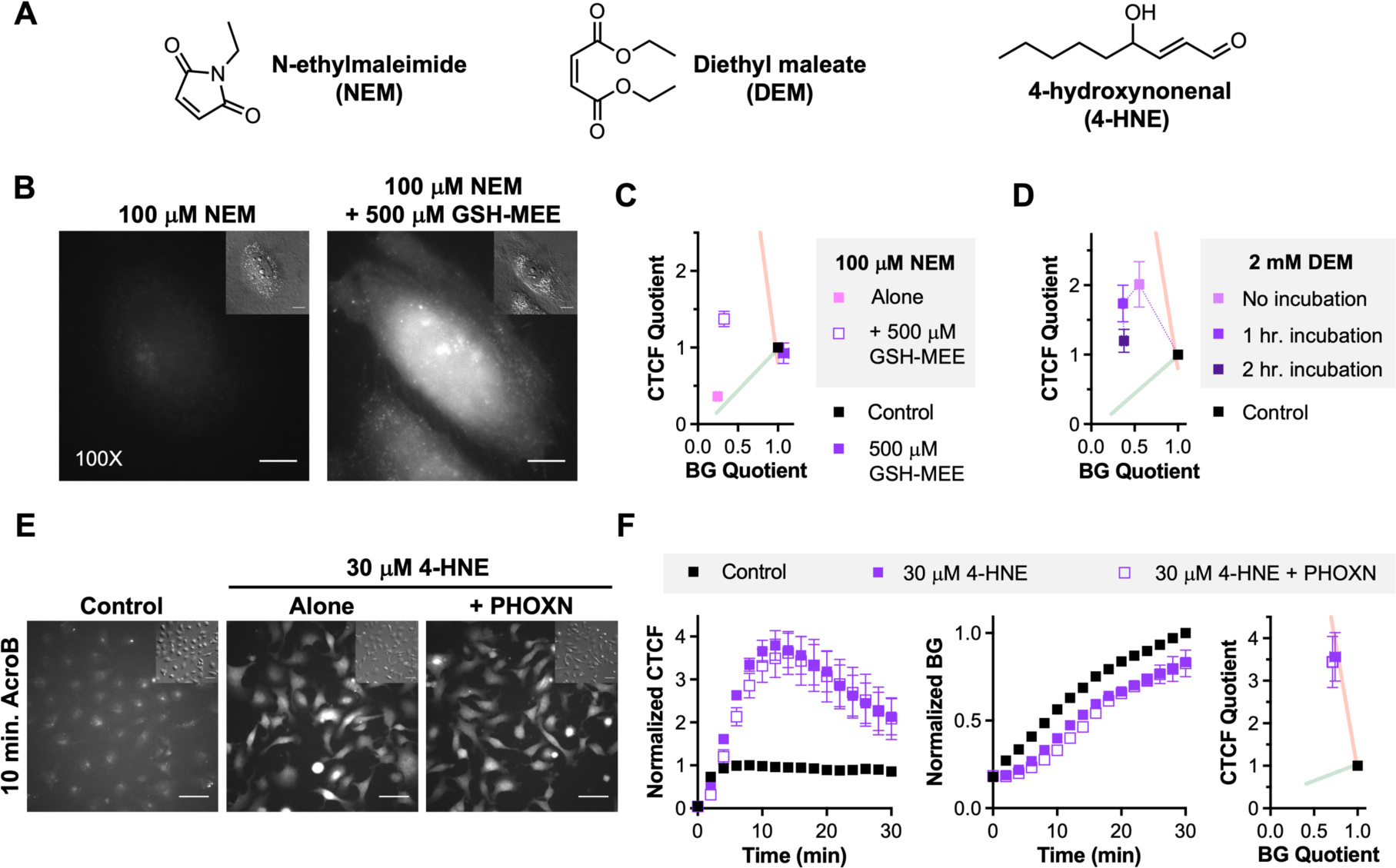
Electrophiles cause LDE-adduct export impairment. **A**, Structures of the electrophiles used: N-ethylmaleimide (NEM), diethyl maleate (DEM) and the LDE 4-hydroxy-2-nonenal (4-HNE). **B**, 100*×* HeLa cell AcroB fluorescence and DIC (inset) images after treatment with 100 μM NEM, 500 μM GSH-MEE and 20-minute AcroB application. GSH-MEE treatment visualizes NEM-caused LDE-adduct export impairment observed as retention of AcroB fluorescent adducts. **C**, ElectrophileQ plot for NEM and/or GSH-MEE treatment conditions presented in **Figure S4A-B**. NEM treatment produces an ElectrophileQ response consistent with GSH conjugation impairment alone; re-introduction of GSH with GSH-MEE treatment visualizes the presence of additional LDE-adduct export impairment. **D**, ElectrophileQ plot for DEM conditions presented in **Figure S4C-D**. No DEM pre-incubation decreases LDE detoxification primarily through LDE-adduct export impairment, while increased incubation times lead to additional GSH conjugation impairment. Line connecting DEM points is a visual aid. **E**, Representative 20*×* AcroB widefield fluorescence and DIC (inset) images of HT1080 cells following treatment with 30 μM 4-HNE alone or following 1 μM PHOXN treatment compared to control conditions. **F**, Normalized CTCF, normalized BG, and ElectrophileQ plots for conditions presented in (**E**). ElectrophileQ plot shows 4-HNE treatment leads to LDE detoxification impairment primarily through LDE-adduct export impairment that is not altered by the presence of the RTA PHOXN. Images taken every 2 min. for 30 min. (20*×*) or every 30 sec. for 30 min. (100*×*) - images taken 20 min. (**B**) or 10 min. (**E**) after AcroB application shown. α_ex_ = 488 nm (0.1 mW for 20*×*, 0.05 mW for 100*×*). Scale bar is 64 μm for 20*×* and 12 μm for 100*×*. AcroB added following the indicated treatment(s). Each data set was normalized to maximum CTCF and BG values obtained using controls performed in parallel. The ElectrophileQ plot analyzes the CTCF and BG values calculated for the images obtained 20 minutes following AcroB application. Sodium orthovanadate and BSO standard curves presented as visual aids. Presented values are average of minimum n = 4 field of view (FOV) for all conditions ± SEM.

A second electrophile we tested was the mildly reactive diethyl maleate (DEM, **Figure 5A**). Diethyl maleate is known to both react with GSH and to alkylate cysteine residues (58, 59). DEM treatment immediately prior to AcroB imaging produced visible retention of AcroB adducts (**Figure S4C**, no incubation condition) and a large increase in CTCF and decrease in BG (**Figure S4D**). Evaluating the LDE detoxification impairment using ElectrophileQ identified mixed inhibition with greater contribution from LDE-adduct export (**Figure 5D**). With increasing lengths of DEM pre-incubation (1 and 2 hrs.), overall fluorescence decreased (**Figure S4C-D**) and the points on the ElectrophileQ plot moved down towards greater contribution of GSH conjugation impairment, demonstrating that DEM induces significant LDE-adduct export inhibition first, followed by GSH depletion.

We subsequently treated HT1080 cells with 4-HNE, the prototypical LDE compound. Incubation of HT1080 cells with 30 μM 4-HNE for 30 minutes prior to AcroB imaging induced retention of fluorescent AcroB adducts (**Figure 5E-F**). The relationship between the increase in CTCF and decrease in BG was consistent with the inhibition of MRP1-mediated LDE-adduct export response observed during ferroptosis as shown by ElectrophileQ (**Figure 5F**). Testing of greater [4-HNE] led to cell death.

In order to demonstrate that LDEs produced by lipid peroxidation are the cause of the LDE adduct export impairment observed during ferroptosis, we treated cells with the lipophilic RTA PHOXN to prevent lipid peroxidation prior to treatment with 4-HNE. In contrast to how RTAs prevented LDE adduct export inhibition following ferroptosis induction (**Figure S2D**) (36) or lipid peroxidation induction (**Figure 4F**), the inhibition of LDE-adduct export caused by 4-HNE was not altered by pre-treatment with PHOXN (**Figure 5E-F**). This is consistent with LDEs acting downstream of PUFA-OOH generation to inhibit LDE-adduct export and isolates LDEs as the mediators of impaired MRP1 activity during ferroptosis.

Both lipid peroxidation and treatment with exogenous electrophiles, including 4-HNE, recapitulate the LDE-adduct export impairment that is a hallmark of ferroptosis. Our results are consistent with and support the known inhibition of MRP channels by LDEs (42–44) and the detected alkylation of MRP1 channels by LDEs during ferroptosis (39). Taken together, these findings indicate that lipid peroxidation during ferroptosis and subsequent breakdown of PUFA-OOH generates high levels of LDEs that inhibit MRP1 channel activity through alkylation.

## Discussion

A growing body of evidence supports a critical role for LDEs in the execution of ferroptosis. In addition to the GSH-dependent LDE detoxification pathway, the aldo-keto reductase (AKR) and aldehyde dehydrogenase (ALDH) enzyme classes metabolize and detoxify LDEs (60–63). Multiple studies have linked decreased AKR and ALDH activity with increased ferroptosis susceptibility, suggesting that successful LDE clearance at least delays ferroptotic cell death (19, 64, 65). While it has long been known that LDE levels are increased during ferroptosis, recent reports showed that sublethal levels of 4-HNE and RSL3 or erastin display synergistic behaviour to induce ferroptotic cell death (65) and position LDEs as mediators of altered metabolism in ferroptotic cells (66, 67). Additionally, treatment with conjugated PUFAs, which can react by peroxyl radical addition mechanisms to more readily form LDEs and truncated PLs compared to non-conjugated PUFAs (22, 68), leads to elevated LDE-mediated damage and induces ferroptotic cell death without the need for exposure to a canonical ferroptosis inducer (69). Widespread protein LDE alkylation (>1000 targets) has been mapped during RSL3-induced ferroptosis, identifying proteins involved in LDE metabolism (incl. ALDHs, an AKR family member, multiple GST isoforms and ABCC1/MRP1), lipid metabolism (incl. MBOAT7 and ACSL4) and cellular antioxidant defense systems (incl. thioredoxin, TXN, and peroxiredoxin, PRDX, family members) (39). The LDE alkylation of proteins specifically implicated in ferroptosis execution was also identified, including those upstream of PUFA-OOH accumulation (incl. VDAC2 and the system x_c_^-^ heavy chain SLC3A2) and those downstream (incl. TRP channels, multiple subunits of the Na^+^/K^+^-ATPase, and Piezo1 channels) (39). However, prior to this work, a demonstration of LDE-mediated altered protein function during ferroptosis was lacking.

Here, we monitored LDE detoxification ability across a panel of FINs using ElectrophileQ. The screening of FIN Types II-IV (RSL3, FIN56 and FINO_2_) showed a consistent mode of LDE detoxification impairment predominantly caused by inhibition of MRP1-mediated LDE-adduct export (**Figure 2**). Testing of a Type I FIN (erastin) with ElectrophileQ revealed mixed inhibition of both the GSH conjugation and LDE-adduct export steps of the LDE detoxification pathway (**Figure 3**). Therefore, with respect to LDE metabolism, we demonstrate that there are two categories of FINs: i) FINs that primarily cause LDE-adduct export impairment once sufficient LDE alkylation has occurred to inhibit MRP1 (ex. RSL3, FIN56, FINO_2_) and ii) FINs that affect LDE detoxification directly by GSH depletion/conjugation impairment and subsequently lead to additional export impairment following progression of ferroptosis (ex. erastin). Importantly, this work demonstrates that regardless of the specific mechanism of ferroptosis induction, the elevated levels of LDEs produced by lipid peroxidation following treatment with each FIN are sufficient to inhibit MRP1-mediated LDE-adduct export (**Figures 2-3**).

Lipid peroxidation induction, with H_2_O_2_ or cumene hydroperoxide, and exogenous 4-HNE treatment each recapitulate the LDE-adduct export impairment that occurs during ferroptosis (**Figure 4-5**). Additionally, in the presence of PHOXN, a potent RTA, 4-HNE treatment still caused inhibition of LDE-adduct export, identifying LDEs produced following lipid peroxidation as the cause of altered MRP1 function during ferroptosis.

This real time visualization of the functional impact of ferroptotic LDE exposure on MRP channels has important implications for our understanding of ferroptotic cell death. Effectively, a protein (MRP1) that was previously shown to be (i) inhibited by LDE alkylation (42–44) and (ii) was identified in the screen of LDE alkylation targets during ferroptosis (39) is now shown in this work to be (iii) functionally altered following LDE exposure during ferroptosis.

Taking a step back, MRP1 channels are just one example of many proteins that are involved in ferroptosis execution and suppression that (i) are known to be activated/inhibited by LDE alkylation and (ii) were shown to be alkylated by LDEs during ferroptosis (39). Our demonstration that MRP1 function is modified by LDEs *during ferroptosis* positions LDE alkylation as the mechanism underlying altered activity of countless proteins during ferroptotic cell death (**Scheme 1**).

**Scheme 1.**
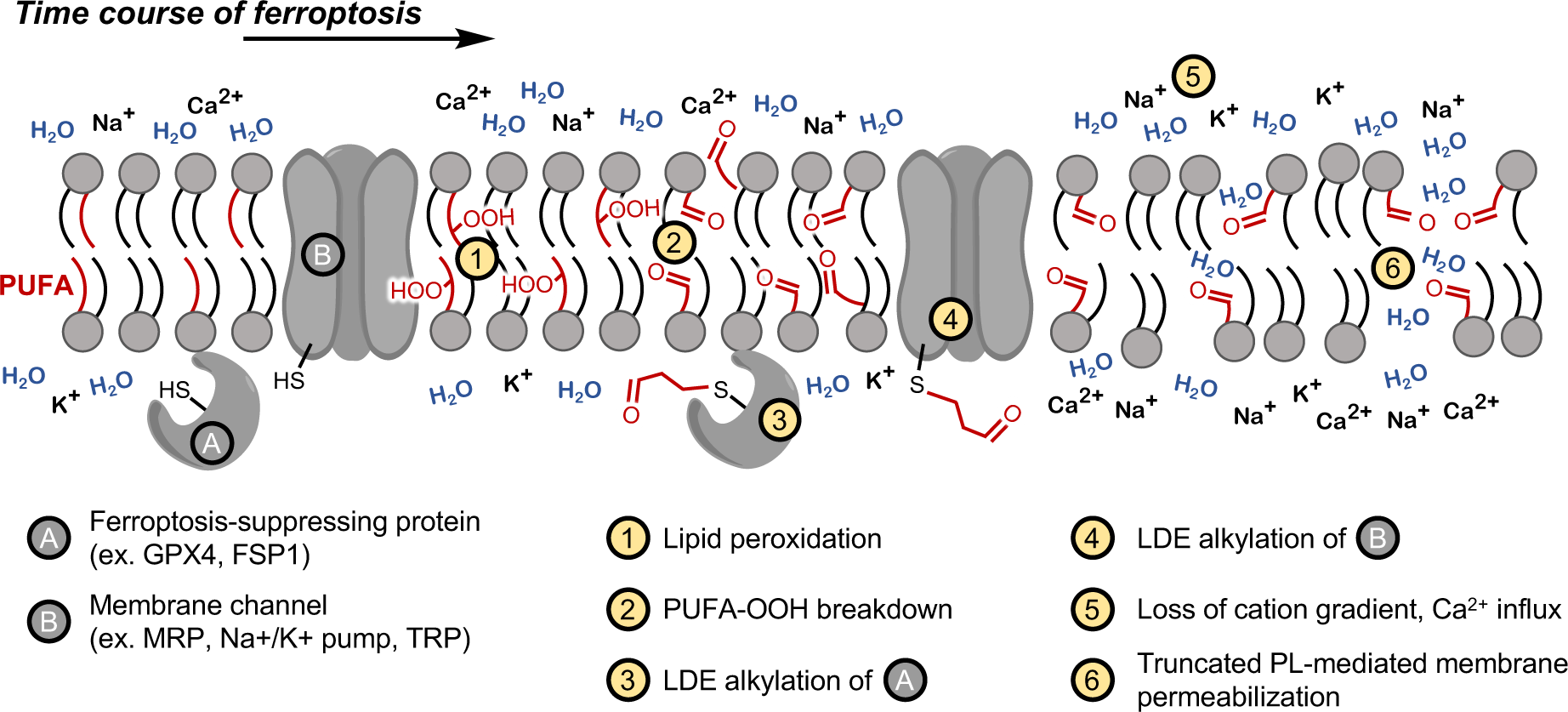
A potential timeline of ferroptotic membrane damage.

For example, inhibition of the Na^+^/K^+^-ATPase by an undetermined mechanism was recently identified during ferroptosis, contributing to critical loss of the cation gradient across the plasma membrane (32). Considering that (i) inactivation of the Na^+^/K^+^-ATPase by LDE alkylation associated with altered cation balance is well characterized (70–72) and (ii) multiple subunits of the Na^+^/K^+^-ATPase were identified in the screen of LDE alkylation targets during ferroptosis (39), we can now posit that the inactivation of the Na^+^/K^+^-ATPase during ferroptosis is caused by LDE alkylation. Similarly, the TRP channels also implicated in the execution of ferroptosis (32, 73–75) can be activated by LDE alkylation (76–78) and multiple TRP family members were identified as ferroptotic LDE targets (39). It remains to be seen if the alkylation of Piezo1 that has been observed during ferroptosis accounts for channel activation (39). The elevated generation and accumulation of LDEs during ferroptosis likely means that alkylation of one single protein will not be identified as the critical step in ferroptotic cell death, but rather the buildup of widespread LDE-mediated altered protein function destroys cell metabolism and integrity.

A potential timeline of membrane damage during ferroptosis is taking shape (**Scheme 1**). In healthy cells, endogenous RTAs and ferroptosis suppressing proteins, such as GPX4 and FSP1, alongside other cellular antioxidant enzymes, such as peroxiredoxin (PRXN) and thioredoxin (TRX), maintain low levels of lipid peroxidation/hydroperoxides. However, ferroptosis begins when homeostasis is lost and lipid peroxidation levels increase (*Step 1*) (4). The accumulating PUFA-OOHs break down to form LDEs and truncated PLs (*Step 2*) (38). LDEs will then alkylate and inactivate a plethora of cysteine-containing antioxidant and aldehyde protective proteins, such as PRDXs, TXNs, AKRs, ALDHs, SLC3A2, and GSTs (39), leading to increased levels of lipid peroxidation and LDE/truncated PL accumulation in a positive feedback loop (*Step 3*). This concept is supported by the indirect inactivation of GPX4 during FINO_2_ induced ferroptosis (17). LDEs also feed forward to alkylate and alter the activity of membrane channels, such as MRP1, TRP channels, Piezo1, and the Na^+^/K^+^-ATPase (*Step 4*), leading to calcium influx and loss of the cation gradient (*Step 5*) (32). Alongside the protein damage caused by LDEs, accumulating truncated PLs lead to eventual membrane permeabilization (*Step 6*) (23, 27).

The breakdown of PUFA-OOHs to form LDEs and truncated PLs may represent a point of no return during ferroptosis and offers a unified explanation of the molecular mechanism of ferroptotic cell death. LDE alkylation can rationalize the modified protein functions currently characterized in late stage ferroptosis, while truncated PLs are sufficient to induce membrane permeabilization independent of the protein pore formation required in other forms of cell death (e.g. NINJ1) (32, 79, 80). The discovery of additional roles for LDEs in the ferroptotic context is likely, as LDEs represent potential mediators of ferroptosis propagation and have been shown to interfere with the immune response (81).

## Conclusion

The molecular mechanism of ferroptotic cell death has been the focus of much research. Here, through a mechanistic investigation of LDE detoxification impairment during ferroptosis, we position LDEs not simply as products of lipid peroxidation, but as executors of damage that leads to ferroptotic cell death. Using the ElectrophileQ assay developed to image GSH-mediated LDE detoxification in live cells, we identified inhibition of MRP-mediated GS-LDE adduct export during ferroptosis induced by a series of FINs (RSL3, FINO_2_, FIN56 and erastin). The induction of cellular lipid peroxidation was shown to replicate this export inhibition, and treatment with 4-HNE, the best characterized LDE, led to impairment of MRP1 function even during inhibition of lipid peroxidation. Our work provides a real-time visualization of LDE-mediated altered protein function during ferroptosis and sets a precedent for a unified mechanism of membrane damage during ferroptosis relying only on lipid hydroperoxide breakdown to truncated PLs and LDEs. This molecular explanation of ferroptotic cell death sets the stage for ferroptosis therapeutic strategies that not only act on lipid peroxidation, but that additionally target the executors of ferroptotic cell death: truncated PLs and LDEs.

## Author Contributions

A.T.M.V.K. and G.C. together conceptualized the project and prepared the manuscript.

A.T.M.V.K. performed the experiments, G.C. directed the research.

## Acknowledgements

1. G. C. is grateful to the Natural Sciences and Engineering Research Council of Canada (NSERC) and the Canadian Foundation for Innovation (CFI) for funding. A.T.M.V.K. is grateful to NSERC and the Fonds de Recherche du Québec – Nature et Technologies (FRQNT) for postgraduate scholarships.

## Supporting Information

Supporting information document contains the Materials and Methods along with Supporting Figures S1-S4.

## Notes

### Competing Interest Statement

The authors have declared no competing interest.

