## Supporting Information for "MRP1 inhibition by lipid-derived electrophiles during ferroptosis illustrates a role for protein alkylation in ferroptotic cell death"

### Contents


- Materials and Methods (Page S3)
  - Materials (Page S3)
  - Cell Culture (Page S3)
  - Imaging (Page S3)
    - Widefield microscopy (Page S3)
    - General imaging conditions (Page S3)
    - Reversan (Page S4)
    - MK571 (Page S4)
    - RSL3 (Page S4)
    - Phenoxazine (Page S4)
    - FIN56 (Page S4)
    - $FINO_2$ (Page S5)
    - Erastin (Page S5)
    - Etoposide (Page S5)
    - Hydrogen peroxide (Page S5)
    - Cumene hydroperoxide and copper (II) sulfate (Page S5)
    - N-ethyl maleimide (Page S5)
    - Diethyl maleate (Page S5)
    - 4-hydroxy-2-nonenal (Page S5)
    - Glutathione monoethyl ester (Page S6)
  - Image Analysis (Page S6)
    - ElectrophileQ assay analysis (Page S6)

- Supporting Figures (Page S7)
  - **Figure S1.** Additional MRP1 inhibitors reversan and MK571 corroborate the ElectrophileQ response of sodium orthovanadate. (Page S7)
  - **Figure S2.** Supporting data to **Figure 2.** (Page S8)
  - **Figure S3.** Lipid peroxidation, not simply cell death, is required to inhibit LDE-adduct export. Supporting data to **Figure 4.** (Page S9)
  - **Figure S4.** Supporting data to **Figure 5.** (Page S11)

- References (Page S12)

### Materials and methods

#### Materials

Buffers, imaging media, growth media and all other reagents for cell culture were purchased through ThermoFisher Scientific. AcroB was synthesized according to previously reported procedures (1). $FINO_2$, erastin and 4-HNE were purchased from Cayman Chemical through Cedarlane Labs. Hydrogen peroxide was purchases through ThermoFisher Scientific. All other chemicals were purchased from Sigma-Aldrich, Co. and used without further purification.

#### Cell culture

HeLa cells (ATCC CCL-2) and HT-1080 cells (CCL-121) were cultured in Dulbecco's modified Eagle's medium (DMEM) containing high glucose, L-glutamine, phenol red, and sodium pyruvate (Gibco), supplemented with 10% fetal bovine serum (FBS, Gibco), 1% penicillin-streptomycin and, for HT-1080 cells, 1% non-essential amino acids (NEAA, Gibco) – hereafter "DMEM" refers to DMEM including the indicated growth supplements. Cells were maintained at 37°C (5% CO2) in a humidified atmosphere. Cells were trypsinized and split 1/10 (HeLa) or 1/20 biweekly (HT-1080) when 90% confluency was reached. Cells were passaged a maximum of 30 times and monitored for mycoplasma infection.

#### Imaging

*Widefield microscopy.* Widefield fluorescence and differential interference contrast (DIC) imaging were performed using a widefield objective-based setup consisting of an inverted microscope (Nikon Eclipse Ti or Nikon Eclipse Ti2) equipped with a Perfect Focus System (PFS) with either a 100× oil-immersion objective (Nikon CFI SR Apochromat TIRF 100×, NA = 1.49) or a 20× air objective (Nikon CFI Plan Apo VC 20× objective, NA = 0.75). Cells were maintained at 37°C and 5% $CO_2$ in a humidified atmosphere using a stage-top incubator (Tokai Hit). 488 nm laser excitation was used at powers of 0.1 mW for 20× and 0.05 mW for 100× imaging measured out of the objective; the beam was coupled into the microscope objective using a multiband beam splitter (ZT488/640rpc, Chroma Technology). Emission was spectrally filtered with a ZET488/640m emission filter (Chroma Technology). A motorized filter block turret was used for multichannel imaging to change between the 488 and DIC channels to obtain a corresponding DIC image with each widefield fluorescence image. Fluorescence emission was collected though the same objective and captured on a back illuminated electron multiplying charge coupled device (EM-CCD) camera (Andor iXon Ultra DU-897). Optical configuration controlled through Nikon NIS-Elements software: auto exposure (300 ms), readout mode (EM Gain 1 MHz at 16-bit), EM gain multiplier (17), conversion gain (Gain 3).

*General imaging conditions.* Unless otherwise indicated, cells were plated 24 hours prior to microscope imaging on a 35 mm glass imaging dish (FluoroDish, World Precision Instruments, Inc.) pre-coated with fibronectin (1 µg/cm2 – for 100× imaging), on an 8-well ibidi-Treat µ-slide (for HeLa 20× imaging) or on fibronectin coated 8-well glass-bottomed ibidi µ-slide (for HT-1080 20× imaging) in DMEM containing growth factors. In order to reach approximately 50% confluency during imaging, 50,000 trypsinized cells suspended in 1 mL DMEM were plated on FluoroDishes and 15,000 trypsinized cells suspended in 300 µL DMEM were plated in each well of the 8-well µ-slides. For imaging, the growth media was removed and replaced with Live Cell Imaging Solution (LCIS, ThermoFisher) – LCIS supplemented with 5 mM glucose (LCIS + glucose) for HT-1080 cells. More specific imaging details for each treatment are below.

After 24-hour incubation on the imaging dish, treatment was performed (see specific conditions below) before cell media was removed and cells were washed 3 times with LCIS, and 1 mL (FluoroDish) or 200 µL (8-well µ-slide) LCIS was left on the cells (LCIS + glucose for HT-1080 cells). Once loaded onto the microscope, cell fields of view (FOV) for imaging were located and their coordinates saved on the PFS; 1 FOV was used for 100× imaging on the FluoroDish, while 2 FOV were chosen in each of the 8 wells on the ibidi µ-slide (16 FOV per imaging dish, see below).

AcroB dye solutions were prepared from a 30 µM stock solution in DMSO. 5 µL of the stock was added to 495 µL LCIS (LCIS + glucose for HT-1080 cells) to obtain a stock solution of 300 nM and a final percentage DMSO on the cells of 0.33%. Immediately before imaging, dye was added as the 300 nM stock solution in LCIS to reach a final concentration in the imaging media of 100 nM (500 µL added to the FluoroDish, 100 µL added to 8-well ibidi µ-slide). The volume of imaging media was set to a uniform 300 µL for ibidi µ-slide experiments as fluorescence background quantification is dependent on the volume of extracellular media; the use of the humidified incubator for imaging also ensures minimal evaporation of imaging media. For 20× widefield imaging, each FOV was imaged every 2 minutes for 30 minutes. For 100× widefield AcroB imaging, cells were imaged on both the DIC and 488 nm fluorescence channels every 30 seconds for 30 minutes. Any deviations from this protocol during the treatments are indicated below.

*Reversan.* 5 mg/mL stock solution (11.32 mM) in DMSO was used and 8.83 µL was diluted into 991 µL DMEM + NEAA to obtain a working solution of 100 µM Reversan. 1/2 and 1/10 dilutions were performed before media exchange onto HT-1080 cells for the 30-minute incubation with 1-10 µM Reversan at 37°C. The respective [Reversan] was present in the LCIS + glucose imaging media following washing.

*MK571.* 5 mg/mL stock solution (9.3 mM) in sterile deionized water was used and 5.37 µL was diluted into 995 µL DMEM + NEAA to obtain a working solution of 50 µM MK571. 1/5 and 1/25 dilutions were performed before media exchange onto HT-1080 cells for the 30-minute incubation with 2-50 µM MK571 at 37°C. The respective [MK571] was present in the LCIS + glucose imaging media following washing.

*RSL3*. 75 µM stock solution of RSL3 in DMSO was used and 16.6 µL was diluted into 484 µL DMEM + NEAA to obtain a working solution of 2.5 µM RSL3. 1/10 dilution onto HT-1080 cells was performed for the 0.5-2-hour incubation with 250 nM RSL3 at 37°C. RSL3 was not present in the LCIS + glucose imaging media following washing.

*Phenoxazine (PHOXN).* 10 mg/mL stock solution of PHOXN was prepared in DMSO and diluted into DMEM + NEAA to a final concentration of 3 µM PHOXN. A 1/3 dilution was performed onto cells to reach a final [PHOXN] of 1 µM for a 30-minute pre-incubation prior to any FIN or electrophile treatment.

*FIN56*. 1 mg/mL stock solution (9.66 mM) in DMSO was used and 2.6 µL was diluted into 497 µL DMEM + NEAA to obtain a working solution of 50 µM FIN56. 1/10 dilution onto HT-1080 cells was performed for the 1–3-hour incubation with 5 µM FIN56 at 37°C. FIN56 was not present in the LCIS + glucose imaging media following washing.

*$FINO_2$.* 1 mg/mL stock solution (3.9 mM) in DMSO was used and 7.7 µL was diluted into 292 µL DMEM + NEAA to obtain a working solution of 100 µM $FINO_2$. 1/10 dilution onto HT-1080 cells was performed for the 1-4–hour incubation with 10 µM $FINO_2$ at 37°C. $FINO_2$ was not present in the LCIS + glucose imaging media following washing.

*Erastin.* 1 mg/mL stock solution (1.83 mM) in DMSO was used and 8.2 µL was diluted into 492 µL DMEM + NEAA to obtain a working solution of 30 µM erastin. 1/3 dilution onto HT-1080 cells was performed for the 9–12-hour incubation with 10 µM erastin at 37°C. Erastin was not present in the LCIS + glucose imaging media following washing.

*Etoposide.* 10 mM stock solution in DMSO was used and 50 µL was diluted into 450 µL DMEM to obtain a working solution of 100 µM etoposide. Media exchange onto HeLa cells was performed for the 24-hour incubation with 100 µM etoposide at 37°C. Etoposide was not present in the LCIS imaging media following washing.

*Hydrogen peroxide.* 1 M stock solution in PBS was used. 20 µL was diluted into 980 µL DMEM to obtain a working solution of 20 mM $H_2O_2$. 1/10 dilution onto HeLa cells was performed for the 2-hour incubation with 2 mM $H_2O_2$ at 37°C. $H_2O_2$ was not present in the LCIS imaging media following washing.

*Cumene hydroperoxide and copper (II) sulfate.* 33 mM cumeneOOH stock solution in PBS was used and 20 µL was diluted into 980 µL DMEM to obtain a 660 µM cumeneOOH working solution. 668 mM copper (II) sulfate stock in sterile distilled water was used and 90 µL was diluted into 910 µL DMEM to obtain a 600 µM copper (II) sulfate working solution. A combined oxidant solution was obtained by adding 450 µL cumeneOOH working solution and 50 µL copper (II) sulfate working solution to obtain 600 µM cumeneOOH and 60 µM copper (II) sulfate. 1/3 dilution onto HeLa cells was performed for the 1-2-hour incubation with 200 µM cumeneOOH and 20 µM copper (II) sulfate or following further 1/2 dilution for treatment with 100 µM cumeneOOH and 10 µM copper (II) sulfate. PMHC pre-treatment was performed starting with a 10 mg/mL PMHC stock (45.4 mM) in DMSO that was diluted to obtain 10 µM and 50 µM PMHC solutions in DMEM for media exchange onto HeLa cells 30 minutes prior to oxidant treatment. CumeneOOH, copper (II) sulfate and PMHC were not present in the LCIS imaging media following washing.

*N-ethyl maleimide (NEM).* 5 mg was dissolved in 400 µL PBS to obtain a 100 mM stock solution. 10 µL was diluted into 990 µL DMEM to obtain a working solution of 1 mM NEM. 1/10 dilution onto HeLa cells was performed for the 30-minute incubation with 100 µM NEM at 37°C. NEM was not present in the LCIS imaging media following washing.

*Diethyl maleate (DEM).* 10 µL neat DEM was added to 990 µL DMSO to obtain a 79.8 µM DEM stock solution. 100.2 µL were diluted into 3.9 mL DMEM to obtain a 2 mM DMEM working solution. Media exchange onto HeLa cells was performed for the 2 hr.-incubation with 2 mM DEM. DEM was not present in the LCIS imaging media following washing.

*4-hydroxy-2-nonenal (4-HNE).* 1 mg/mL stock solution (6.4 mM) in DMSO was used and 23.4 µL was diluted into 477 µL DMEM + NEAA to obtain a working solution of 300 µM 4-HNE. 1/10 dilution onto HT-1080 cells was performed for the 30-minute incubation with 30 μM 4-HNE at 37°C. 4-HNE was not present in the LCIS + glucose imaging media following washing.

*Glutathione monoethyl ester (GSH-MEE).* 5 mg/mL stock solution (30 mM) stock solution in PBS was used and 16.77 μL was diluted into 1 mL DMEM + NEAA to obtain a working solution of 500 μM GSH-MEE. Media exchange onto cells was performed for the 30-minute incubation with 500 μM GSH-MEE at 37°C. GSH-MEE was not present in the LCIS + glucose imaging media following washing. Treatment with GSH-MEE was done following completion of the incubation time with erastin or 4-HNE prior to imaging.

**Image analysis**

All images were processed using the FIJI imaging processing package (2). Widefield fluorescence corrected total cell fluorescence (CTCF) and background fluorescence (BG) were quantified as previously described (3, 4).

Normalization of the CTCF and BG values was performed by division of the calculated values for a given condition by the maximum calculated value for control conditions performed in parallel. CTCF and BG values are presented normalized to controls performed in parallel to facilitate comparison between different conditions and incubation lengths.

*ElectrophileQ assay analysis.* All values for the ElectrophileQ plots are calculated using CTCF and BG intensity values obtained from quantification of 20× widefield imaging experiments. Values chosen for comparison are those calculated from the images taken 20 minutes after AcroB application. CTCF quotient is calculated by taking the quotient of the average CTCF from the given treatment condition and the average control CTCF value. BG quotient is calculated by taking the quotient of the average fluorescence background from the given treatment condition and the average control fluorescence background value. Due to the use of widefield illumination, photobleaching level is assumed to be uniform between the cell and background regions and is therefore not accounted for in the analysis.

### Supporting Figures

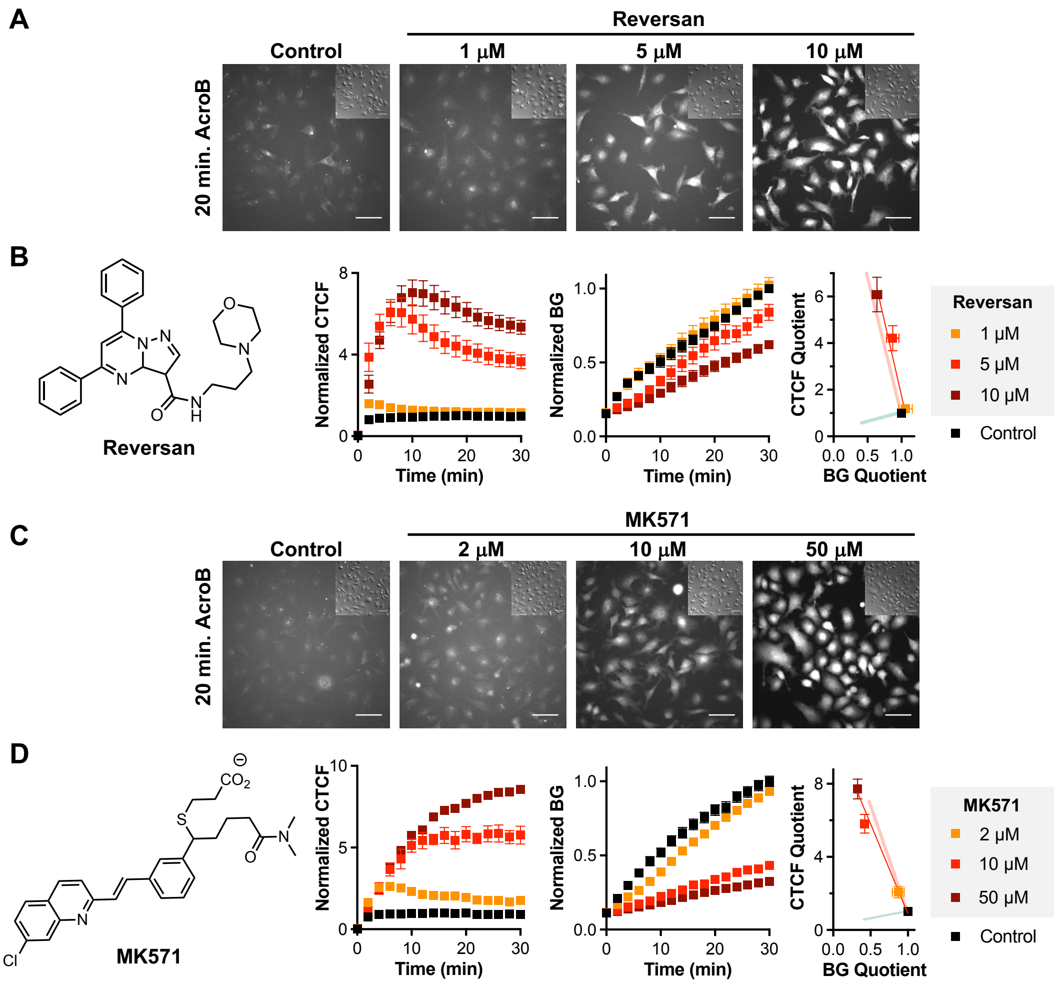


**Figure S1. Additional MRP1 inhibitors reversan and MK571 corroborate the ElectrophileQ response of sodium orthovanadate.** **A**, Representative 20× AcroB widefield fluorescence and DIC (inset) images of HT1080 cells following treatment with increasing concentrations of reversan compared to control conditions. **B**, Chemical structure of Reversan, normalized average CTCF, normalized average BG, and ElectrophileQ plots corresponding to the conditions presented in (**A**). Reversan treatment produced LDE-adduct export that closely matched sodium orthovanadate on the ElectrophileQ plot. **C**, Representative 20× AcroB widefield fluorescence and DIC (inset) images of HT1080 cells following treatment with increasing concentrations of MK571 compared to control conditions. **D**, Chemical structure of MK571, normalized average CTCF, normalized average BG, and ElectrophileQ plots corresponding to the conditions presented in (**C**). MK571 treatment produced LDE-adduct export that closely matched sodium orthovanadate on the ElectrophileQ plot. AcroB was added following the indicated treatment. Each data set was normalized to maximum CTCF and BG values obtained using controls performed in parallel. The ElectrophileQ plot analyzes the CTCF and BG values calculated for the images obtained 20 minutes following AcroB application. Sodium orthovanadate and BSO standard curves presented as visual aids. Presented values are average of minimum n = 4 field of view (FOV) for all conditions ± SEM.

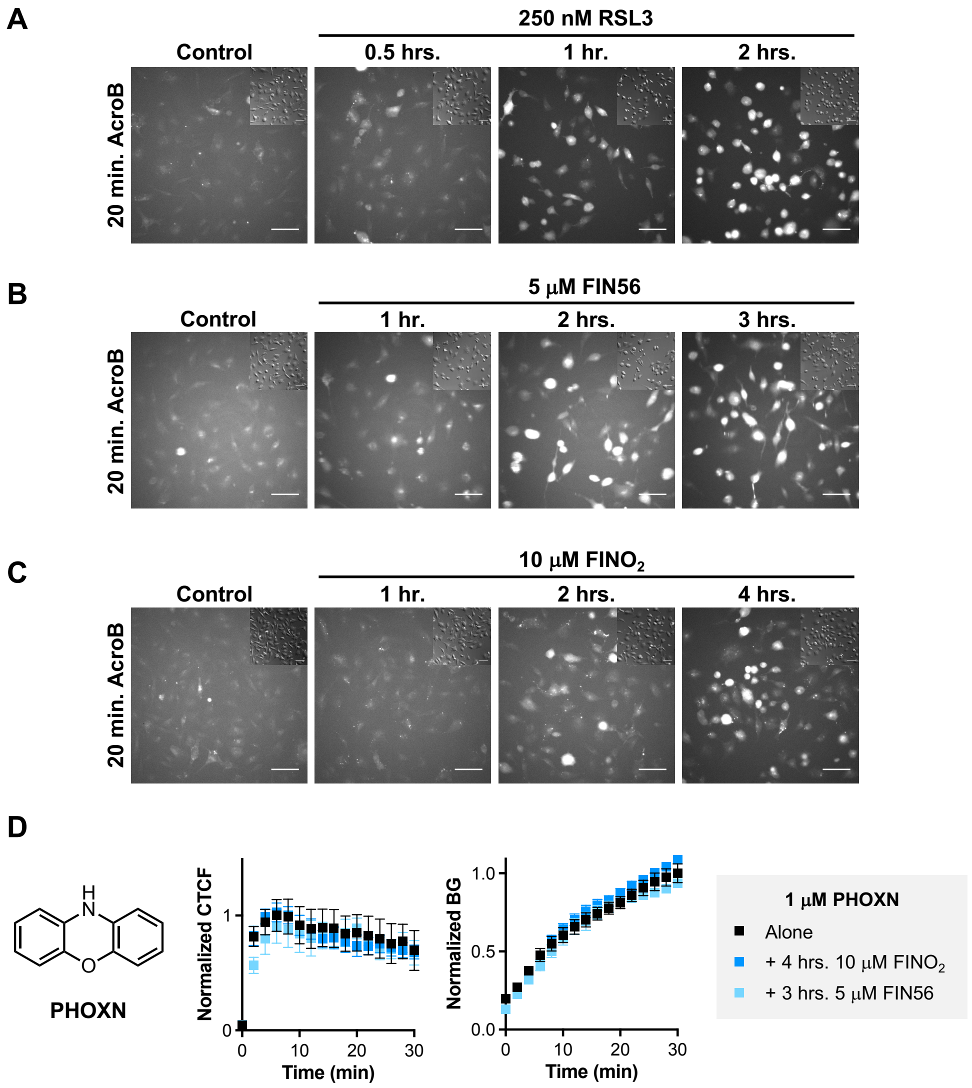


**Figure S2. Supporting data to Figure 2. A-C**, Representative 20× AcroB widefield fluorescence and DIC (inset) images of HT1080 cells following treatment with (**A**) RSL3, (**B**) FIN56 or (**C**) $FINO_2$ for varying incubation times show consistent increase in cell fluorescence and decrease in background fluorescence compared to control. Images were taken every 2 minutes for 30 minutes following application with 100 nM AcroB (image 20 minutes following AcroB application shown). Scale bar is 64 µm. $\lambda_{ex}$ = 488 nm (0.1 mW). **D**, Chemical structure of PHOXN, normalized CTCF and BG curves for $FINO_2$ and FIN56 treatment following pre-treatment with 1 µM PHOXN obtained from 20x images show that the presence of an RTA prevents altered LDE detoxification due to these FINs. Presented values are average of minimum n = 4 field of view (FOV) for all conditions ± SEM.

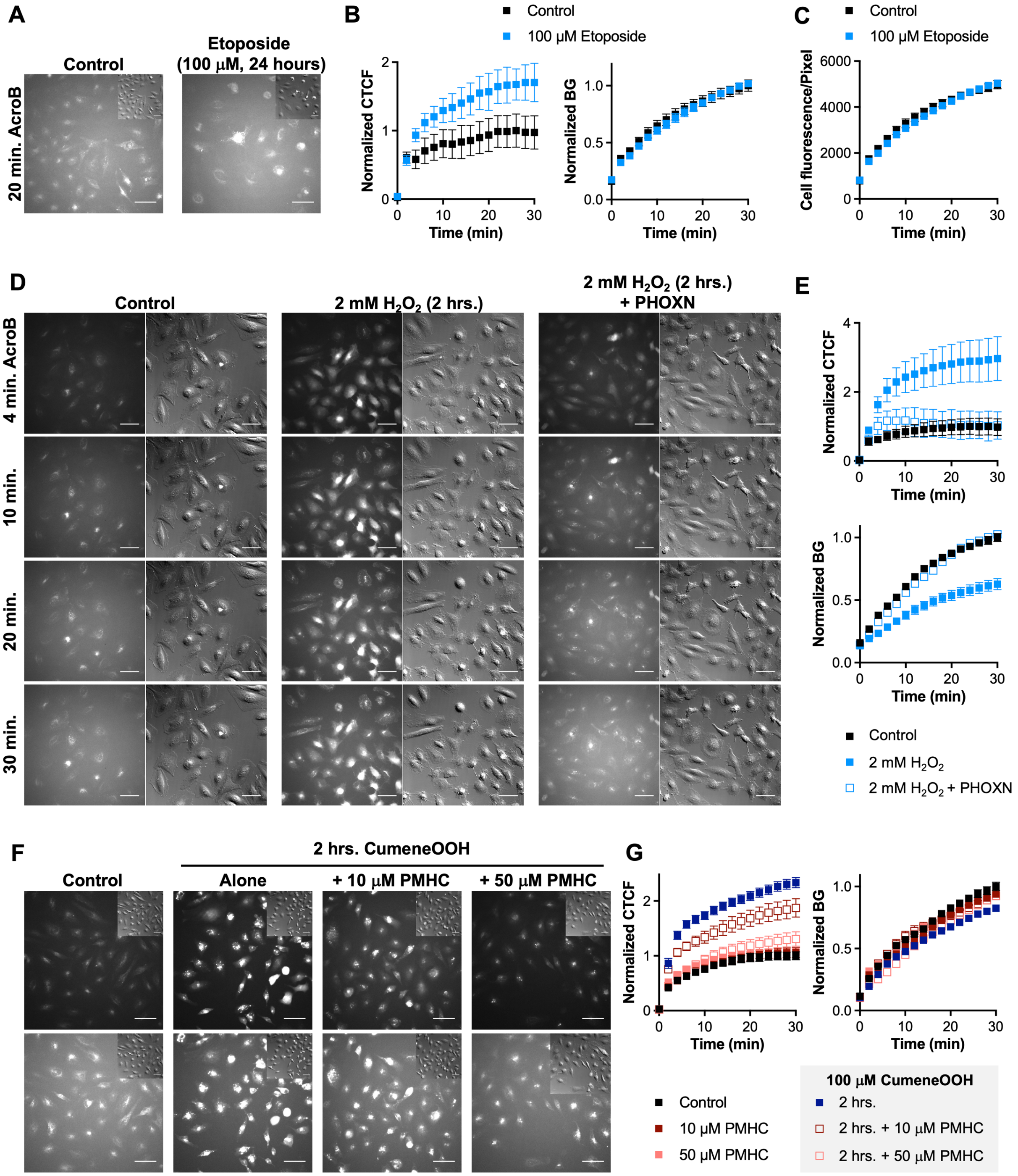
A
Control
Etoposide (100 µM, 24 hours)
20 min. AcroB
B
Control
100 µM Etoposide
Normalized CTCF
Normalized BG
Time (min)
C
Control
100 µM Etoposide
Cell fluorescence/Pixel
Time (min)
D
Control
2 mM H2O2 (2 hrs.)
2 mM H2O2 (2 hrs.) + PHOXN
4 min. AcroB
10 min.
20 min.
30 min.
E
Normalized CTCF
Normalized BG
Time (min)
Control
2 mM H2O2
2 mM H2O2 + PHOXN
F
Control
2 hrs. CumeneOOH
Alone
+ 10 µM PMHC
+ 50 µM PMHC
G
Normalized CTCF
Normalized BG
Time (min)
Control
10 µM PMHC
50 µM PMHC
100 µM CumeneOOH
2 hrs.
2 hrs. + 10 µM PMHC
2 hrs. + 50 µM PMHC

**Figure S3. Lipid peroxidation, not simply cell death, is required to inhibit LDE-adduct export. Supporting data to Figure 4. A**, Representative 20× AcroB widefield fluorescence and DIC (inset) images of HeLa cells following treatment with 100 μM etoposide for 24 hrs compared to control conditions. **B**, Normalized average CTCF and BG plots corresponding to the conditions presented in (**A**) appear to show increased CTCF vs. control. **C**, Presentation of cell fluorescence per pixel for conditions presented in (**A**) clarifies that total cell fluorescence per unit area has not changed, but rather the etoposide-mediated prevention of cell division and subsequent cell growth accounts for differences observed in (**B**). **D**, Representative 20× AcroB widefield fluorescence (left) and DIC (right) images of HeLa cells following treatment with $H_2O_2$ and PHOXN compared to control conditions. Images at multiple time points following AcroB application are shown to highlight that while PHOXN pre-treatment recovers response on the ElectrophileQ plot following $H_2O_2$ exposure (**Figure 4B**), morphological changes are still observed despite PHOXN pre-treatment (DIC images). **E**, Normalized average CTCF and BG plots corresponding to the conditions presented in (**D**). **F**, Representative 20× AcroB widefield fluorescence and DIC (inset) images of HeLa cells following indicated treatments with 100 μM cumeneOOH, 10 μM copper (II) sulfate and PMHC compared to control conditions. **G**, Normalized average CTCF and BG plots corresponding to the conditions presented in (**F**). Corresponding ElectrophileQ plot presented in **Figure 4F**. AcroB was added following the indicated treatment. Each data set was normalized to maximum CTCF and BG values obtained using controls performed in parallel. Presented values are average of minimum n = 4 field of view (FOV) for all conditions ± SEM.

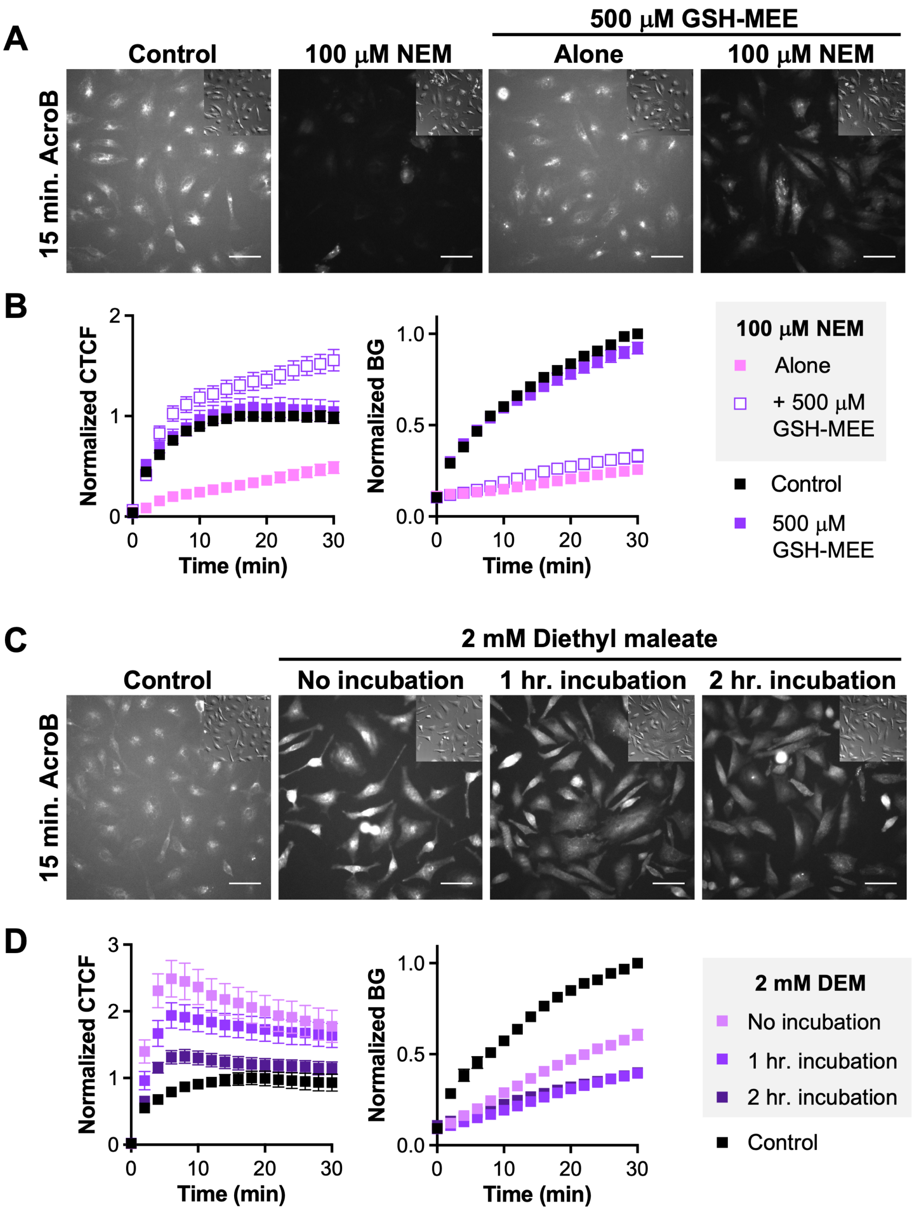


**Figure S4. Supporting data to Figure 5. A**, Representative 20× AcroB widefield fluorescence and DIC (inset) images of HeLa cells following treatment with NEM and/or GSH-MEE compared to control conditions. **B**, Normalized average CTCF and BG plots corresponding to the conditions presented in (**A**). Corresponding ElectrophileQ plot presented in **Figure 5C**. **C**, Representative 20× AcroB widefield fluorescence and DIC (inset) images of HeLa cells following treatment with increasing incubation times of DEM compared to control conditions. **D**, Normalized average CTCF and BG plots corresponding to the conditions presented in (**C**). Corresponding ElectrophileQ plot presented in **Figure 5D**. AcroB was added following the indicated treatment. Each data set was normalized to maximum CTCF and BG values obtained using controls performed in parallel. Presented values are average of minimum n = 4 field of view (FOV) for all conditions ± SEM.
